# IgA displays site- and subclass-specific glycoform differences despite equal glycoenzyme expression

**DOI:** 10.1101/2024.12.11.627887

**Authors:** David Falck, Maria V. Sokolova, Carolien A.M. Koeleman, Vanessa Irumva, Philipp Kirchner, Sebastian R. Schulz, Katja G. Schmidt, Thomas Harrer, Arif B. Ekici, Bernd Spriewald, Georg Schett, Manfred Wuhrer, Martin Herrmann, Ulrike Steffen

## Abstract

**Background:** Glycosylation is an important posttranslational modification of proteins and in most cases indispensable for proper protein function. Like most soluble proteins, IgA, the second most prevalent antibody in human serum, contains several N- and O-glycosylation sites. While for IgG the impact of Fc glycosylation on effector functions and inflammatory potential has been intensively studied, only little is known for IgA. In addition, only glimpses exist regarding the regulation of IgA glycosylation. We have previously shown that IgA1 and IgA2 differ functionally and also show differences in their glycosylation pattern. The more pro-inflammatory IgA2 which is linked to autoimmune diseases displays decreased sialylation, galactosylation, fucosylation and bisection as compared to IgA1. In the present study, we aimed to investigate these differences in glycosylation in detail and to explore the mechanisms underlying them.

**Methods:** IgA1 and IgA2 was isolated from serum of 12 healthy donors. Site specific glycosylation was analyzed by mass spectrometry. In addition, human bone marrow plasma cells were investigated using single cell mRNA sequencing, flow cytometry and ELISpot.

**Results:** We found that certain glycoforms greatly differ in their abundance between IgA1 and IgA2 while others are equally abundant. Overall, the IgA2 glycans displayed a more immature phenotype with a higher prevalence of oligomannose and fewer fully processed glycans. Of note, these differences can’t be explained by differences in the glycosylation enzyme machinery as mRNA sequencing and flow cytometry analysis showed equal enzyme expression in IgA1 and IgA2 producing plasma cells. ELISpot analysis suggested a slightly increased antibody production rate in IgA2 producing plasma cells which might contribute to its lower glycan processing rates. But this difference was only minor, suggesting that further factors such as steric accessibility determine glycan processing. This is supported by the fact that glycans at different positions on the same IgA chain differ dramatically in fucosylation, sialylation and bisection.

**Conclusion:** In summary, our detailed overview of IgA1 and IgA2 glycosylation shows a class, subclass, and site-specific glycosylation fingerprint, most likely due to structural differences of the protein backbones.

## Background

Glycosylation is a very common and complex posttranslational modification of proteins. In humans, the most prevalent glycosylation forms are 1) N-glycosylation in which glycans are attached to asparagine residues embedded into the consensus motive sequence Asn-X-Ser (with X being any amino acid except for proline) and 2) O-glycosylation in which glycans are attached to serine or threonine (or less often tyrosine or other hydroxyl-containing residues). In addition, less common glycosylation forms, such as C-glycosylation (attached to tryptophan residues) and S-glycosylation (attached to cysteine residues) have been described [1]. Especially extracellular proteins contain one or multiple glycosylation sites, important for protein conformation, function, localization and half-life. Glycosylation strongly enhances the variability of proteins due to alterations in the presence of glycans and oligosaccharide composition, referred to as macro- and microheterogeneity, respectively [2].

A very prominent and well-defined example for the impact of glycosylation on protein function is immunoglobulin G (IgG). IgG contains one conserved N-glycosylation site on asparagine 297 located in the CH2 domain of the heavy chain. The attached biantennary complex-type glycan reaches into the pocket between the two heavy chains and, therefore, affects the Fc domain conformation and the affinity of IgG to Fcγ receptors (FcγRs). Changes in galactosylation, sialylation and fucosylation are known to alter IgG effector functions and have been associated with various diseases, including rheumatoid arthritis, systemic lupus erythematosus, and coronavirus disease 2019 (COVID-19) [3-7]. In addition, pharmacological industry employs glycoengineering to improve half-life and effector functions of therapeutic antibodies [8].

In humans, IgA is the most frequently produced Ig and with about 2-3 mg/ml second most abundant Ig in the circulation [9]. Humans have functional genes for IgA1 and IgA2. In contrast to IgG with only one conserved N-glycosylation site, IgA1 harbors two and IgA2 even four (or five in case of IgA2m(2) allotype) (Fig. 1) [9]. As in IgG, the attached glycans are typically of the biantennary complex type comprising a core of four N-acetylglucosamine and three mannose residues (= core heptamer) to which additional sugars, such as core fucose, bisecting N-acetylglucosamine, galactose and sialic acid at one or both arms can be attached. In addition, in contrast to IgG, early biosynthetic pathway forms like oligomannose and hybrid type glycans are found in IgA in minor quantities. In addition, the elongated hinge region of IgA1 harbors six O-glycosylation sites of which between four to five are occupied on average [10]. The impact of these glycans on IgA effector function remains largely unknown. Non-secreted serum IgA is recognized by Fcα receptor I (FcαRI) mainly expressed on myeloid cells, such as neutrophils, macrophages, or some subsets of dendritic cells [11]. Although it seems that the IgA glycans do not interfere with binding of IgA to the FcαRI, they can influence the immune system [12]. Terminal sialic acid at the complex N-glycans at the C-terminal tailpiece of IgA has been shown to interact with influenza viruses increasing their neutralization [13]. Abnormal O-glycosylation is mainly associated with kidney disease in IgA nephropathy and IgA vasculitis. Galactose-deficient IgA1 found in patients with IgA nephropathy binds mannose-binding lectin and thereby activates complement via the lectin pathway [14, 15]. In addition, changes in O- and N-glycosylation patterns have been observed in several autoimmune diseases such as systemic lupus erythematosus, rheumatoid arthritis and primary Sjögren’s syndrome [16-20].

**Fig. 1:**
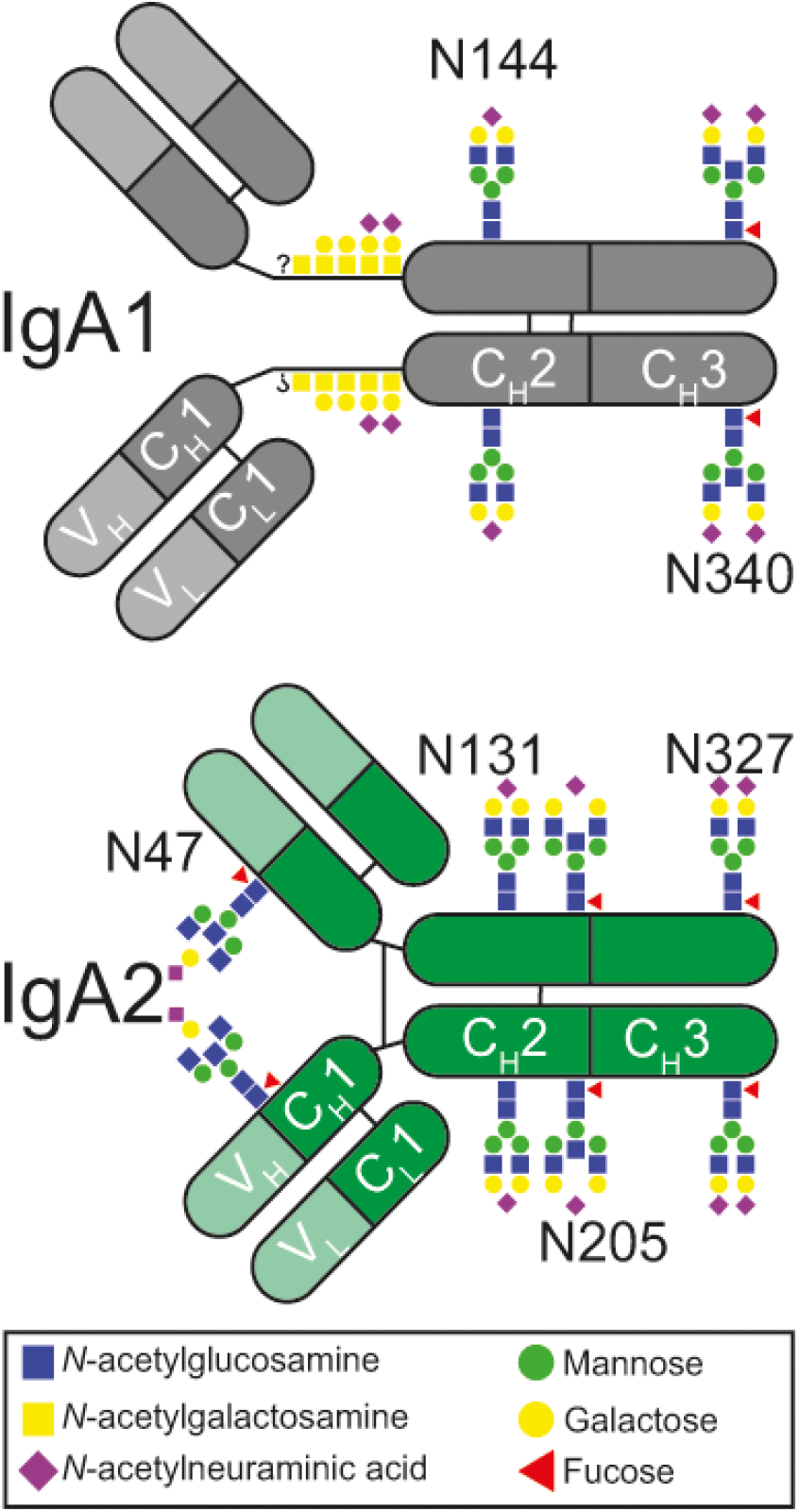
Schematic illustration of IgA N- and O-glycosylation sites.

By binding to FcαRI, IgA can induce either inflammatory or anti-inflammatory effects. Comparing the two subclasses it has been shown that IgA2 elicits stronger pro-inflammatory responses than IgA1 [21, 22]. Interestingly, IgA1 and IgA2 differ in their glycosylation with IgA1 showing in general higher degrees of sialylation, galactosylation, fucosylation and bisecting N-acetylglucosamine. We found that sialic acid removal from IgA1 increases its inflammatory properties to levels comparable to IgA2, suggesting that glycosylation regulates IgA function [21].

The regulation of IgA glycosylation is not well understood. Protein glycosylation is not template-driven, but determined by the composition of the glycosylation machinery and the accessibility of the modified protein [23]. N-glycosylation already starts during the translation process in the endoplasmic reticulum with the attachment of an initial mannose-rich oligosaccharide structure by the enzyme oligosaccharyltransferase to the growing polypeptide chain. After protein folding, the terminal glucose and mannose residues are removed in the endoplasmic reticulum. This process continues in the Golgi apparatus before the truncated oligomannose structure is again extended by glycosyltransferases. This comprises the initiation of the antennae with *N*-acetylglucosamine and the further attachment of galactose, sialic acid, core fucose and/ or bisecting *N*-acetylglucosamine [23]. Regarding IgG glycosylation, it is assumed that the abundance of glycoenzymes determines the degree of glycan processing (e.g. the amount of galactosylation, sialylation or fucosylation). Indeed, genetic and epigenetic variations targeting glycosyltransferases are associated to differences in glycosylation [24-27]. However, the question remains if such variations are also responsible for glycosylation differences between antibody classes and subclasses or if these differences are solely determined by the protein structure.

To shed light on this question, we compared the detailed glycoform composition of IgA1 and IgA2. In addition, we compared glycosyltransferase expression and the antibody production rate in IgA1 and IgA2 expressing plasma cells using single cell sequencing, flow cytometry and ELISpot.

## Methods

### IgA isolation

IgA was isolated from sera of 12 healthy donors (age 40–62 years; 75% female) as described previously [21]. Briefly, the sera were mixed with equal amounts of phosphate-buffered saline (PBS). Total IgA was isolated using a column with peptide M agarose (Invivogen) according to the manufacturer’s instructions. After a pre-elution step with 0.2 M glycine buffer (pH 5.0) to remove unspecifically bound proteins, IgA was eluted with 0.1 M glycine buffer (pH 2.7), immediately neutralized using 1M Tris buffer (pH 9), and dialyzed against PBS (pH 7.2). Then IgA1 and IgA2 were separated using a column with jacalin agarose (Thermo Scientific) according to the manufacturer’s instructions. IgA2 stayed in the flow through. IgA1 bound to the jacalin agarose and was eluted with 0.1 M galactose buffer. Both IgA fractions were concentrated and buffered to PBS using Amicon Ultra Centrifugal Filters (Merck). Purity was tested with western blot analysis using antibodies against IgA1 and IgA2 (Southern biotech).

### Site-specific mass spectrometry glycosylation analysis

An equivalent of 20 μg of IgA1 or IgA2 was dried and dissolved in 10 μl of 100 mM Tris buffer (pH 8.5) containing 1% sodium deoxycholate, 10 mM Tris(2-carboxyethyl)phosphine, and 40 mM chloroacetamide. Denaturation, reduction, and alkylation was performed for 5 min at 95 °C. After cooling, 500 ng of sequencing grade trypsin (Promega) in 50 μl ammonium bicarbonate buffer (pH 8.5) were added and the samples were incubated overnight at 37 °C. Afterward, sodium dodecyl sulfate (SDS) was precipitated with 2 % formic acid. Liquid chromatography–mass spectrometry analysis of the tryptic glycopeptides was performed as described previously [28]. Briefly, (glyco-)peptides were separated using an Acclaim PepMap C18 column (particle size 2 μm, pore size 100 Å, 75 × 150 mm, Thermo Scientific) and analyzed on a quadrupole-time-of-flight-mass spectrometer (Impact HD, Bruker Daltonics) coupled via a Captive Sprayer nano-electrospray ionization source (Bruker Daltonics). Data were processed with LaCyTools version 1.1.0-alpha, build 20181102b [29].

### Single cell mRNA and VDJ sequencing

For the single cell sequencing, cryopreserved human bone marrow mononuclear cells from three healthy donors were purchased from STEMCELL^TM^ Technologies (5-50 Million cells per donor; viability > 93 %; two male African-Americans, 30 and 40 years old; one female Caucasian, 36 years old). Cells were stained for viability with Zombie-Yellow™ Fixable Viability Kit (1:600) for 20 minutes at room temperature; and afterwards with anti-CD14-BV421, anti-CD16-BV421, anti-CD56-BV421, anti-CD38-APC, anti-CD45-FITC, and anti-CD138-PE/Cy7 (all Biolegend). After the staining, plasma cells (CD45^+^CD14^-^CD16^-^CD56^-^CD38^++^CD138^++^) were sorted on a FACS Aria machine. At the output, 1600-3100 live cells were subjected to 10x Chromium Single Cell sequencing library generation with 10X Chromium 5’ GEX/VDJ kits according to the manufacturer’s instructions. GEX Library sequencing was performed on an Illumina HiSeq 2500 sequencer to a depth of about 50 million reads each. In all samples, the mean number of reads per cell was greater than 30000. VDJ Libraries were sequenced to a depth of about 30 million reads resulting in 1000 to 10000 read pairs per cell. Reads were converted to FASTQ format using mkfastq from Cell Ranger 3.1.0 (10X Genomics). Reads were then aligned to the human reference genome provided by 10X Genomics (GEX: GRCh38-3.0.0; VDJ: vdj_GRCh38_alts_ensembl-3.1.0-3.1.0). Alignment was performed with the count or vdj command from Cell Ranger 3.1.0 (10X Genomics). Further QC filtering was performed in the R package Seurat (v.3.1.4) for samples individually. IgH-Annotation was imported from VDJ sequencing results and the 3 samples were merged for SCTransform normalization. The SCTransform normalization included the number of genes per cell and the sample identity as covariates. Standard dimensionality reduction (PCA) was used, with removal of ribosomal, mitochondrial, immunoglobulin and cell-cycle genes from VariableFeatures. UMAP clustering was performed using the Seurat (v.3.1.4) package for R.

### Flow cytometry analysis

Human bone marrow mononuclear cells (1,5 Million cells per stain) were first incubated with Fc Receptor blocking solution Human TruStain FcX™ (Biolegend) and then stained extracellularly for 20 minutes at 4°C with anti-CD38-APC-antibody, anti-CD138-PE/Cy7, anti-CD45-APC/Cy7 (all Biologend), anti-IgA1-PE, anti-IgA2-FITC (both SouthernBiotech). After fixation with eBioscience™ IC Fixation Buffer in the dark at room temperature for 20 minutes, cells were washed with 1X eBioscience™ Permeabilization Buffer and stained intracellularly with anti-Man1A2 or anti-B4GALT1 (both Invitrogen) over night at 4°C. On the following day, secondary anti-rabbit IgG-BV421 (BD) was added for 2 hours at 4°C. Flow cytometry was performed on a Gallios cytofluorometer and evaluated using Kaluza Analysis 2.1 software (both Beckman Coulter).

### Enzyme-linked immunosorbent spot (ELISpot)

ELISpot was performed on 96-well Filtration Multiscreen IP Sterile Plates (Merck). Plates were activated with 70% MeoH, coated with goat F(ab)2 anti-human IgA (Southern Biotec) and incubated overnight at 4°C. Then, the plates were washed with PBS and blocked at least 1 hour at 37°C with PBS supplemented with 2 % fetal calf serum (FCS). Mononuclear bone marrow cells were seeded at 1 million cells/ml and incubated for 14 hours at 37°C and 5 % CO_2_. After discarding the supernatant, plates were washed with PBS and incubated for 1 hour with mouse anti-human IgA1-horse radish peroxidase (HRP) or mouse anti-human IgA2-HRP (both Southern Biotec). Subsequently, plates were washed with PBS/0.05 % Tween20. Plates were developed with 3,3′,5,5′-Tetramethylbenzidine (TMB) substrate for 10-15 min. The reaction was stopped with distilled water and the plates were left at room temperature for drying. The read-out was done on the AID ELISpot Reader, and analysis of the spot size was done with FiJi and GraphPad Prism 9 softwares.

### Statistical analysis

Statistical analysis was performed with GraphPad Prism 9.0.2. Prior testing, all data sets were tested for normality with Shapiro-Wilk test. For comparison of groups with normal distribution, paired two-tailed Student’s t-test was employed. In case of non-normal distribution, groups were compared using Wilcoxon matched-pairs signed rank test. Differences were considered statistically significant when the p-value was less than 0.05. Data are presented as pie charts, scatter plots with mean ± standard error of mean (s.e.m.) or violin plots with medians and inter-quartile ranges.

## Results

### The glycans of the IgA heavy chain strongly differ between glycosylation sites

To compare the glycoforms of IgA1 and IgA2, we analyzed mass spectrometry-derived site-specific glycosylation data from IgA1 and IgA2, isolated separately from the serum of 12 healthy donors[21]. This analysis revealed major differences between the glycoforms of the different glycosylation sites on the IgA heavy chain (Fig. 2A and Supplementary Table 1). The strongest difference can be seen in fucosylation which is almost absent at N144/131, in contrast to prevalences between 80% and 100% at all other N-glycosylation sites (Supplementary Fig. 1) as reported before [12, 30]. Similarly, although less strong, the bisection rate is reduced about 30-50% at N144/131 compared to the other glycosylation sites. The presence of galactose and terminal sialic acid, on the other hand, shows a slightly different picture and increases uniformly from N-terminal to C-terminal sites. These data indicate differences in regulation for the addition of core fucose and bisecting N-acetylglucosamine as compared to the addition of the more distant galactose and sialic acid.

**Fig. 2:**
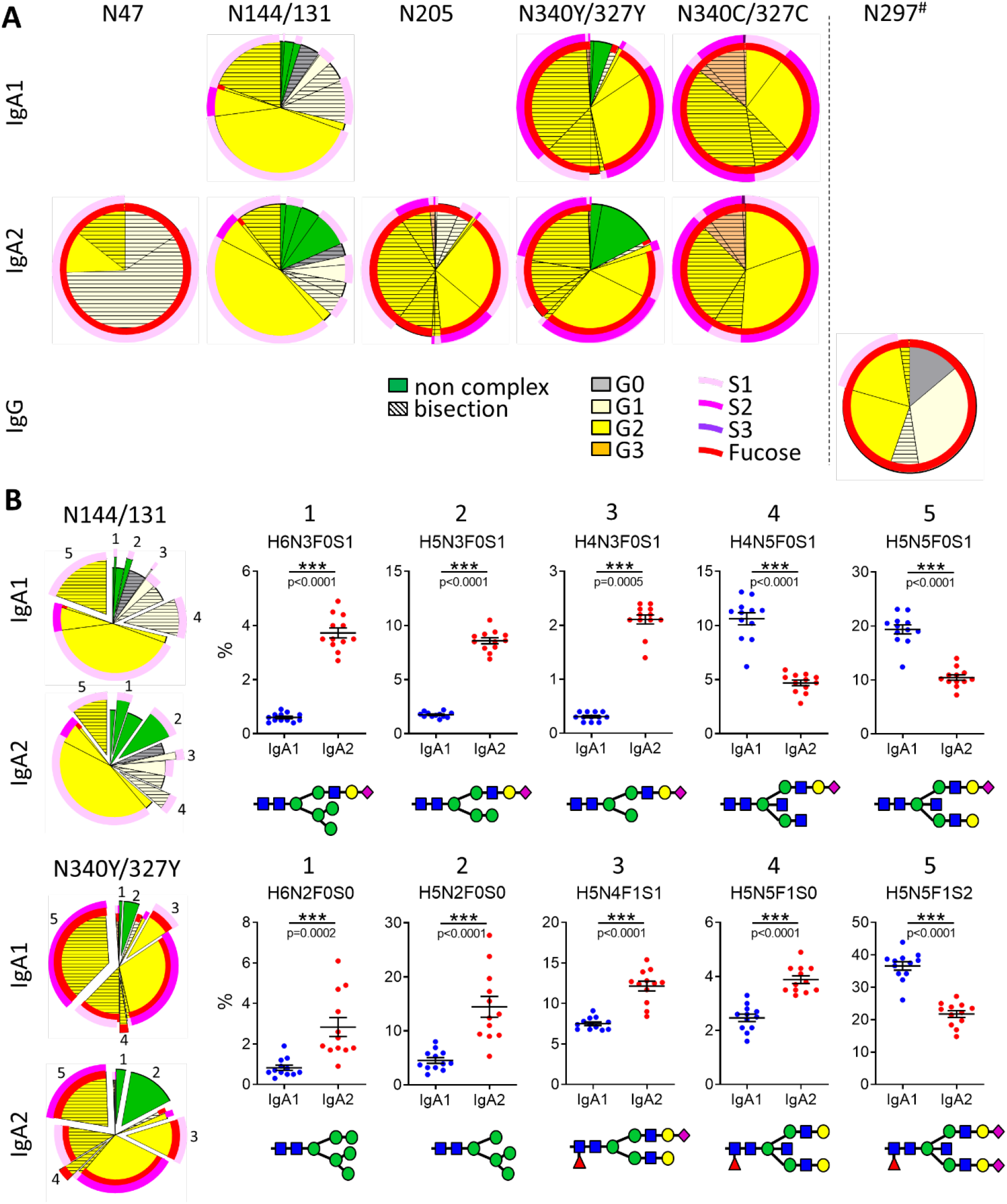
Major differences between the glycoforms at distinct glycosylation sites of IgA1 and IgA2. (**A**) Pie charts of the relative abundances of glycoforms decorating IgA1 and IgA2 glycosylation sites (IgG for reference). Shown are the means of 12 donors. N340/327C and N340/327Y refer to the truncated peptide, lacking terminal tyrosine, and the full peptide, respectively. ^#^ For the IgG glycosylation site, the conventional amino acid number according to literature has been used (e.g. [32]), while glycosylation sites of IgA1 and 2 are indicated according to UniProt numbering (P01876-1 and P01877-1). The presence of fucose is depicted as inner red circles; the degree of sialylation is depicted as outer pink circles. (**B**) Percentage and structure of the 5 glycoforms that differ most in their abundance between IgA1 and IgA2 at the conserved glycosylation sites. Significance was tested with paired t-test. *** = p<0.001; glycan code: H = hexose (mannose or galactose); N = N- acetylglucosamine; F = fucose; S = sialic acid.

The peptide containing the N340/327 glycosylation site exists in a full-length and a truncated form, the latter lacking terminal tyrosine. The biological relevance of these two forms so far remains elusive. Interestingly, these two variants showed differences in their glycan composition with the full-length variant generally having less elaborated glycans than the truncated form (Fig. 2A). For example, the full-length form displays higher amounts of “immature” oligomannose glycoforms while the truncated form displays higher amounts of fully processed tri-antennary glycans.

We made use of small contaminations of IgG in the IgA2 samples to compare the glycan composition of IgA with the Fc N-glycosylation site of IgG. In concordance with the literature, IgG shows much less galactosylation, sialylation and bisection than IgA, while the degree of fucosylation is similar to most IgA glycans (Fig. 2A) [31].

The molecular structures of the glycoforms found in IgA1, IgA2 and IgG are depicted in Supplementary Fig. 2.

### IgA1 and IgA2 are differentially glycosylated

When comparing the glycan composition of the two IgA subclasses, we found that the abundance of some glycoforms strongly differs between IgA1 and IgA2 while others are equally abundant (Fig. 2B and Supplementary Table 1). For both conserved glycosylation sites, IgA1 shows in general more processed glycoforms such as the fully processed H5N5F1S2. Especially, bisection is more prevalent in IgA1. In contrast, IgA2 has a relatively high proportion of “immature” glycoforms that are of non-complex type and still contain outer-arm mannose residues.

### IgA1 and IgA2 plasma cells display similar glycoenzyme expression

To investigate whether the differences in relative glycoform abundances between IgA1 and IgA2 are caused by a different composition of the glycoenzyme machinery in the respective plasma cells, we performed single cell sequencing of bone marrow plasma cells from healthy donors. Of note, all plasma cells clustered well together with a relatively even distribution of all antibody subclasses, suggesting no major differences between IgA1 and IgA2 producing plasma cells (Fig. 3A and Supplementary Fig. 3 for other isotypes). This was supported by the fact that enzymes involved in antibody glycosylation were similarly expressed in IgA1 and IgA2 producing plasma cells (Fig. 3B). Beta-1,4-mannosyl-glycoprotein 4-beta-N-acetylglucosaminyltransferase (MGAT3), responsible for introduction of bisecting N-acetylglucosamine, was unfortunately below the detection limit and could therefore not be analyzed. To validate expression data of glycosylation enzymes important for mannose trimming and galactosylation on protein level, we performed intracellular flow cytometry analyses for mannosidase alpha class 1A member 2 (MAN1A2) and beta-1,4-galactosyltransferase 1 (B4GALT1), respectively [33] [34] (Fig. 3C). In addition, we stained for surface 2,6-sialic acid as a surrogate for beta-galactoside alpha-2,6-sialyltransferase 1 (ST6GAL1) expression, as commercially available antibodies against ST6GAL1 reportedly showed unspecific binding [35]. As already expected from the mRNA sequencing data, we did not see any difference in the amount of these enzymes between IgA1 and IgA2 producing plasma cells (Fig. 3D and Supplementary Fig. 4 for gating strategy). Together these data show that the analyzed glycoenzymes in IgA1 and IgA2 producing plasma cells are similarly expressed.

**Fig. 3:**
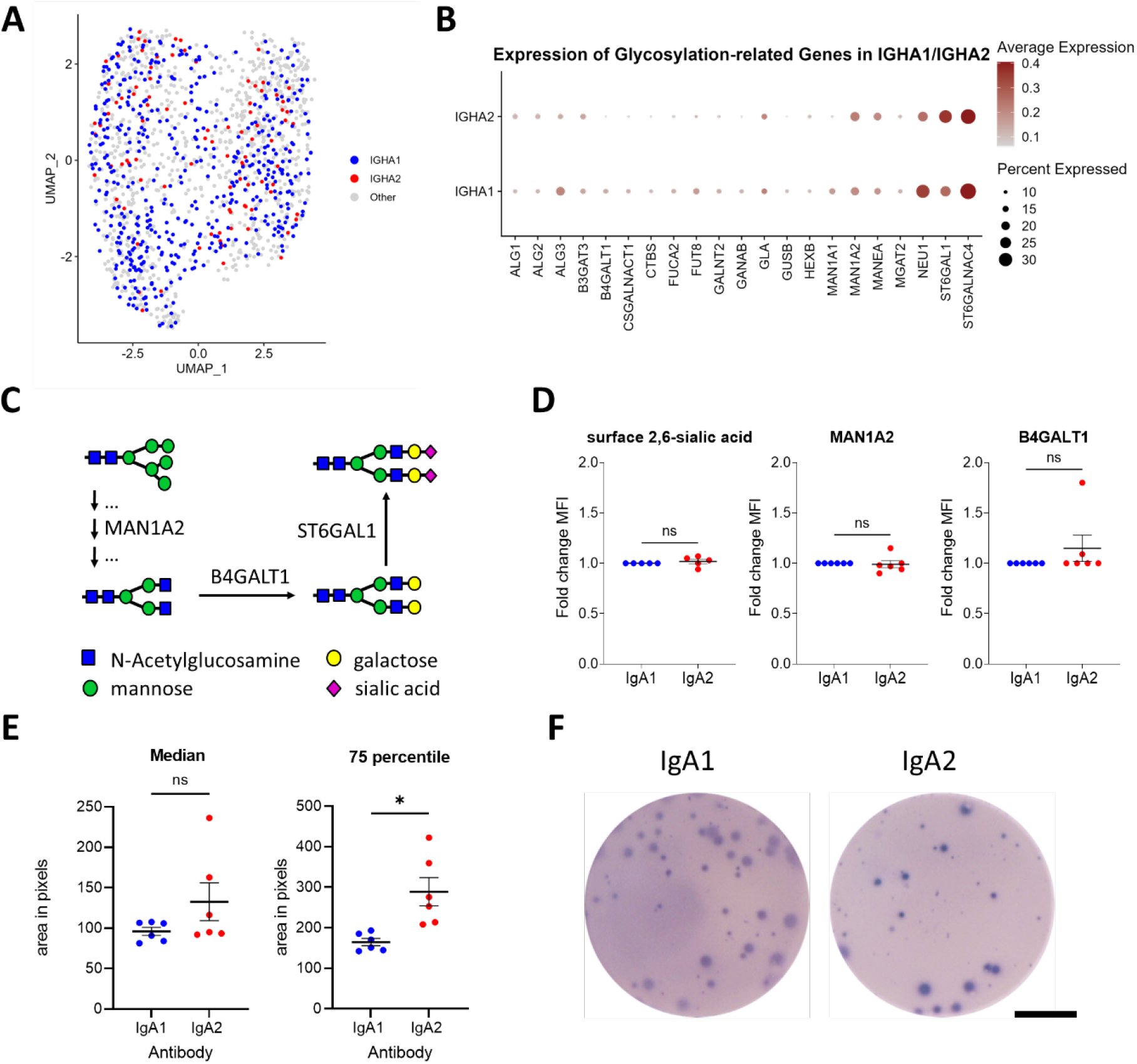
IgA1 and IgA2 producing plasma cells show equal glycosyltransferase amounts, but different antibody release rates. (**A**) Single cell RNA sequencing analysis UMAP representation of plasma cells sorted from bone marrow of healthy donors. Shown are combined data from 3 donors. Heavy chain subclass information was extracted from VDJ sequencing annotations. Here, only IGHA1/2 are highlighted. Other IGHC isotypes can be seen in Supplementary Fig. 3. (**B**) Dot plot for expression of glycosyltransferases in IgHA1 and IgHA2 expressing plasma cells. Dot colors represent mean expression of the genes in each cell group and dot sizes indicate the percentage of cells expressing the respective genes. (**C**) Schematic overview of the contribution of selected enzymes to glycan processing. MAN1A2 is involved in trimming of the outer-arm mannose residues, B4GALT1 and ST6GAL1 add galactose and terminal sialic acid residues, respectively. (**D**) Flow cytometry analysis of IgA1 and IgA2 producing bone marrow plasma cells. Shown is the mean fluorescence intensity (MFI) for surface 2,6-sialic acid (measured by binding of sambuccus nigra agglutinine) and intracellular MAN1A2 and B4GALT1 (measured by binding of specific antibodies). Data are normalized on the MFI of IgA1 producing plasma cells. Every dot represents one donor. N=5-6. **E)** ELISpot analysis of IgA1 and IgA2 producing bone marrow plasma cells. Shown is the median and the 75 percentile of the spot size. Significances were tested with Wilcoxon rank test. * – p<0.05; ns – not significant. (**F**) Representative ELISpot images. Scale bar = 2 mm.

### Plasma cells release slightly more IgA2 than IgA1

As the glycoenzyme to antibody ratio can also be affected by the amount of the produced antibodies, we next investigated IgA1 and IgA2 production by bone marrow plasma cells employing ELISpot. We observed that IgA2 producing plasma cells made around one third larger spots (nearly doubled size if only investigating the quartile of spots) suggesting an increased antibody production rate of these cells (Fig. 3E,F and Supplementary Fig. 5).

## Discussion

Antibody effector functions are strongly influenced by their Fc glycan composition. This has been well established for IgG, and seems also to be the case for IgA [21, 36]. However, the regulation of antibody glycosylation is still unclear.

We found strong differences between the glycan composition of IgA and IgG and, in addition, between the two conserved N-glycosylation sites that are common in IgA1 and IgA2. These IgG and IgA glycosylation profiles qualitatively and quantitatively match previous reports [37, 38]. In general, IgA N-glycans are more extensively galactosylated and sialylated and show a higher rate of bisection than IgG. This might be explained by the fact that the IgG Fc glycan is buried within the pocket of the two IgG heavy chains while the IgA Fc glycans reach out and are therefore easier accessible [32, 39]. This would also explain the fact that the most accessible glycosylation site in IgA, the C-terminal N340/327, contained the glycans with the highest degree on galactosylation and sialylation. However, several findings suggest that in the details the relative activities of different glycotransferases follow more complex laws. For example, the IgG and IgA N47 sites show complete core fucosylation, despite low galactosylation and sialylation, while the IgA glycan at N144/131 mostly lacks fucose, but has high galactosylation. Nonetheless, the high degree of bisection and the recent observation of triantennary structures (personal communication) on IgA N47 are in line with a generally greater accessibility of IgA N47 than IgA N144/131. Also, Thaysen-Andersen and Packer suggested that core fucosylation, maybe due to its proximity to the peptide backbone, is partially determined by hydrophobicity of the protein backbone sequence close to the glycosylation site, while distant sugar modifications are rather determined by protein folding accessibility [40]. Finally, the issue is further complicated by circulating glycosidases that may lead to a lower abundance at accessible sites, most prominently described for neuraminidases [41]. This could be a further factor that contributes to the differences of IgA and IgG glycosylation, as IgG with its significantly longer plasma half-life would be exposed to these circulating glycosidases far more extensively.

Comparing the two IgA subclasses, IgA2 contains more “immature” glycans characterized by a higher amount of mannose residues and the lack of the typical complex structures. While differences between distinct N-glycosylation sites within one IgA molecule can only be explained by steric effects and protein conformation, glycosylation differences between the IgA subclasses could also result from differences in the glycoenzyme composition of the respective plasma cells or the antibody production rate. However, we did not see any differences in glycoenzyme expression on mRNA and protein level between IgA1 and IgA2 producing plasma cells. When we compared their antibody production rate, IgA2 producing plasma cells secreted more IgA. As the addition of sugar residues to proteins is an inefficient process with interactions of the target proteins and glycoenzymes on a stochastic basis, an increased IgA to enzyme ratio could partly explain the lower processing levels of IgA2 glycans. In addition, IgA2 has twice as many N-glycosylation sites as IgA1 and, therefore, also a lower enzyme to glycan ratio. Next to competition for enzymes, competition for nucleotide sugar donors might also play a role. Opposingly, IgA1 contains several O-glycans that are absent in IgA2 and which might compete for nucleotide sugar donors as well. Still, steric differences between IgA1 and IgA2 cannot be excluded as cause for the differences in the glycan composition. Although the amino acid sequence close to the N-glycosylation sites is similar between IgA1 and IgA2, more distant structural differences could affect glycosylation especially at the more central N144/131 glycosylation site. In addition, differences in the levels of the sugar nucleotide donors, thus roughly the metabolic state of the plasma cell, and putative differences in the allocation of glycoenzymes to the antibody-containing Golgi vesicles might have an impact [42].

Regarding the conserved N-glycosylation sites of IgA1 and IgA2, we previously published that IgA2 glycans are less galactosylated, sialylated, bisected, fucosylated and complex than IgA1 [21]. Our detailed analysis here shows that in particular some glycoforms are changed while the proportions of other glycoforms remain unchanged. This might be of importance as different glycan modifications have been shown to interfere with each other. For example, IgG galactosylation further enhances FcγRIIIa affinity of non-fucosylated IgG by twofold [43]. The information about specific changes in glycosylation together with changes in effector functions might, therefore, help to better understand glycosylation regulation and to improve targeted glycoengineering approaches.

## Conclusions

Our study has some limitations. Although our single cell sequencing analysis showed a generally good coverage of glycosylation enzymes, MGAT3, which is important for bisection, escaped detection. In addition, due to the technology it is not possible to know which glycans of the different sites are connected in an individual Igα chain. Lastly, we did not investigate the steric effects on the glycan composition of the individual sites. The strength of our data is the high resolution of glycan composition at distinct glycosylation sites in the two IgA subclasses.

Together, our data show that there are specific differences in the glycan composition both between distinct N-glycosylation sites within one IgA subclass and between the two IgA subclasses at conserved N-glycosylation sites. These differences were independent of the glycoenzyme composition of plasma cells and are therefore most likely caused by structural differences and possibly influenced by the IgA production rates in plasma cells.

## Abbreviations

B4GALT1: beta-1,4-galactosyltransferase 1
COVID-19: coronavirus disease 2019
FcαRI: Fcα receptor I
FcγRs: Fcγ receptors
FCS: fetal calf serum
HRP: horse radish peroxidase
Ig: immunoglobulin
IGHC: immunoglobulin heavy chain
MAN1A2: mannosidase alpha class 1A member 2
MGAT3: beta-1,4-mannosyl-glycoprotein 4-beta-N-acetylglucosaminyltransferase
MFI: mean fluorescence intensity
PBS: phosphate-buffered saline
SDS: sodium dodecyl sulfate
s.e.m.: standard error of mean
ST6GAL1: beta-galactoside alpha-2,6-sialyltransferase 1
TMB: 3,3′,5,5′-Tetramethylbenzidine

## Supplementary Information

Supplementary Table 1

Supplementary Fig. 1

Supplementary Fig. 2

Supplementary Fig. 3

Supplementary Fig. 4

Supplementary Fig. 5

## Declarations

### Ethics approval and consent to participate

All analyses were performed in accordance to the Declaration of Helsinki principles, the institutional guidelines and with the approval of the Ethical Committee of the University Clinic of Erlangen (Permit 277_17 B). All individuals provided informed consent prior to participation in the study.

## Consent for publication

not applicable

## Availability of data and materials

The single cell mRNA sequencing datasets generated and analysed during the current study are available in the GEO repository under the accession number GSE282272, https://www.ncbi.nlm.nih.gov/geo/query/acc.cgi?acc=GSE282272.

## Authors’ contributions

MVS, DF, MH, and US designed the study. MVS, CAMK and VI performed experiments and analyzed the data. PK, SRS and ABE analyzed the sequencing data. BS, GS, and MW gave intellectual input.

MVS, DF, MH, and US wrote the initial manuscript. All authors read, corrected and approved the manuscript.

## Acknowledgement

We thank Sandra Loskarn for excellent technical assistance and Uwe Appelt and Markus Mroz for the cell sort.

