## supplementary material for "IgA displays site- and subclass-specific glycoform differences despite equal glycoenzyme expression"

**Supplementary Table 1: Glycoform distribution in IgA1, IgA2 and IgG.** Shown are mean frequencies (in %) of 12 donors of individual glycoforms at distinct N-glycosylation sites of IgA1 and IgA2 as well as IgG. N340C/327C refers to the truncated peptide lacking terminal tyrosine. # For the IgG glycosylation site, the conventional amino acid number according to literature has been used, while glycosylation sites of IgA1 and 2 are indicated according to UniProt numbering. Glycan code: H = hexose (mannose or galactose); N = N-acetylglucosamine; F = fucose; S = sialic acid.

| Glycoform | IgA2 N47 | IgA1 N144 | IgA2 N131 | IgA2 N205 | IgA1 N340Y | IgA2 N327Y | IgA1 N340C | IgA2 N327C | IgG N297# |
|---|---|---|---|---|---|---|---|---|---|
| **H3N4F1S0** | - | - | - | - | - | - | - | - | 13.9 |
| **H3N5F0S0** | - | 4.8 | 3.1 | - | - | - | - | - | - |
| **H4N3F0S1** | - | 0.3 | 2.1 | - | - | - | - | - | - |
| **H4N4F0S1** | - | 3.2 | 4.3 | - | - | - | - | - | - |
| **H4N4F1S0** | - | - | - | - | - | - | - | - | 33.7 |
| **H4N4F1S1** | - | - | - | 0.4 | - | - | - | - | - |
| **H4N5F0S0** | - | 4.8 | 3.7 | - | - | - | - | - | - |
| **H4N5F0S1** | - | 10.6 | 4.7 | - | - | - | - | - | - |
| **H4N5F1S0** | 15.4 | - | - | 5.8 | 1.7 | 1.2 | - | - | 7.7 |
| **H4N5F1S1** | 59.0 | - | - | 3.5 | - | - | - | - | - |
| **H5N2F0S0** | - | 2.3 | 4.7 | - | 4.5 | 14.4 | - | - | - |
| **H5N3F0S1** | - | 1.8 | 8.6 | - | - | - | - | - | - |
| **H5N4F0S0** | - | 1.8 | 2.4 | - | - | - | - | - | - |
| **H5N4F0S1** | - | 42.3 | 43.8 | 0.7 | - | - | - | - | - |
| **H5N4F0S2** | - | 7.0 | 6.1 | 0.4 | 1.0 | 1.7 | - | - | - |
| **H5N4F1S0** | - | - | - | - | - | - | - | - | 24.2 |
| **H5N4F1S1** | 11.2 | 0.8 | 0.9 | 25.3 | 7.5 | 12.1 | 10.3 | 19.5 | 18.0 |
| **H5N4F1S2** | - | - | - | 12.5 | 30.8 | 29.0 | 27.3 | 31.7 | - |
| **H5N5F0S1** | - | 19.3 | 10.4 | 1.7 | 1.1 | 1.3 | - | - | - |
| **H5N5F0S2** | - | - | - | 0.6 | - | - | - | - | - |
| **H5N5F1S0** | - | - | - | 9.4 | 2.5 | 3.9 | - | - | 2.6 |
| **H5N5F1S1** | 14.4 | - | - | 30.2 | 12.4 | 11.2 | 10.0 | 7.6 | - |
| **H5N5F1S2** | - | - | - | 8.0 | 36.6 | 21.7 | 38.0 | 28.8 | - |
| **H6N2F0S0** | - | 0.4 | 1.4 | - | 0.8 | 2.8 | - | - | - |
| **H6N3F0S1** | - | 0.6 | 3.7 | - | - | - | - | - | - |
| **H6N5F1S1** | - | - | - | 1.2 | 0.7 | 0.3 | 2.4 | 2.5 | - |
| **H6N5F1S2** | - | - | - | 0.2 | 0.4 | 0.3 | 11.3 | 9.1 | - |
| **H6N5F1S3** | - | - | - | - | - | - | 0.7 | 0.8 | - |

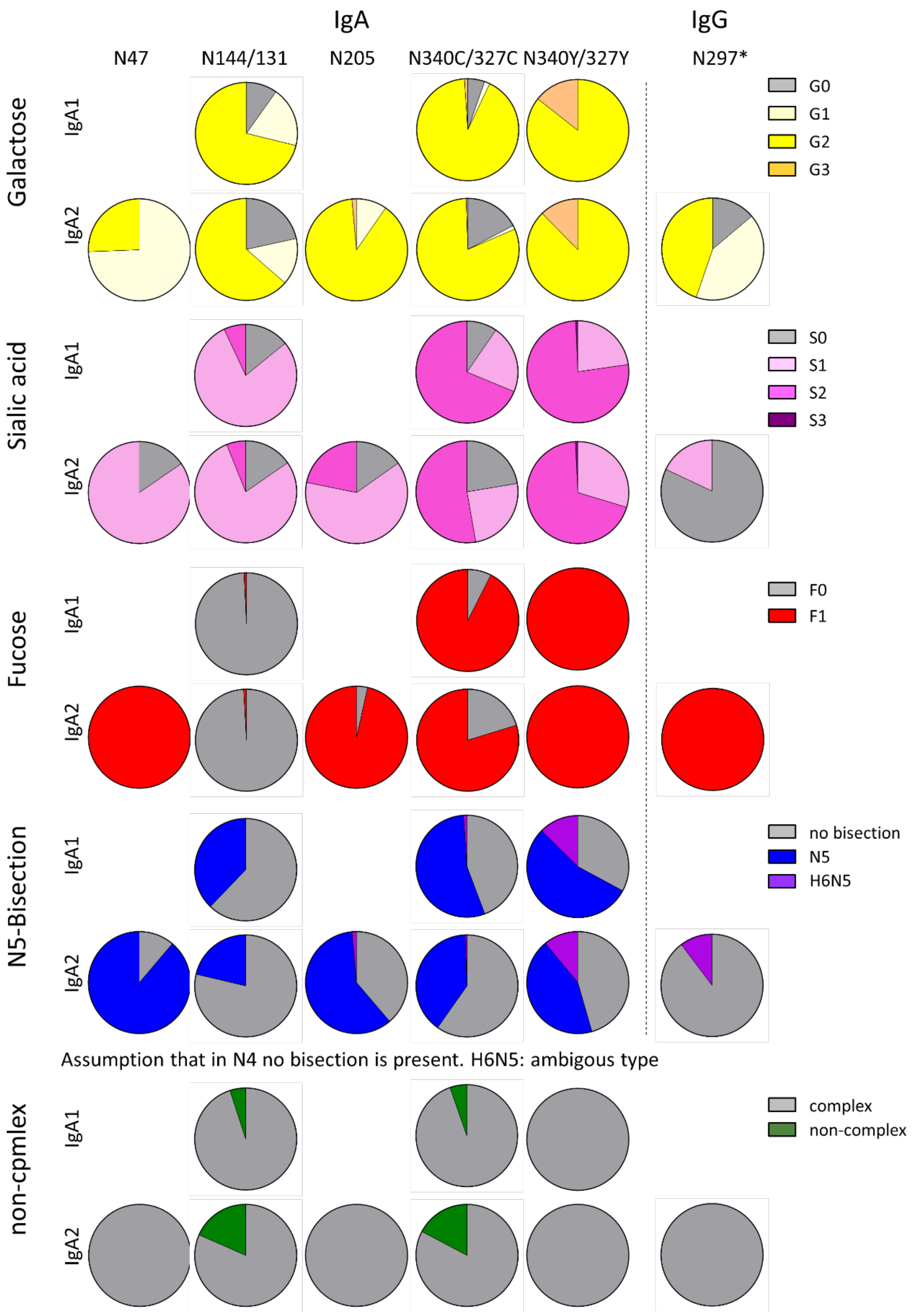


**Supplementary Fig. 1: Glycan trait distribution at distinct glycosylation sites of IgA1 and IgA2.** Pie charts of the frequencies of glycan traits at the depicted glycosylation sites of IgA1 and IgA2 as well as IgG. Shown are mean frequencies of 12 donors. N340C/327C refers to the truncated peptide lacking terminal tyrosine. * For the IgG glycosylation site, the conventional amino acid number according to literature has been used, while glycosylation sites of IgA1 and 2 are indicated according to UniProt numbering.

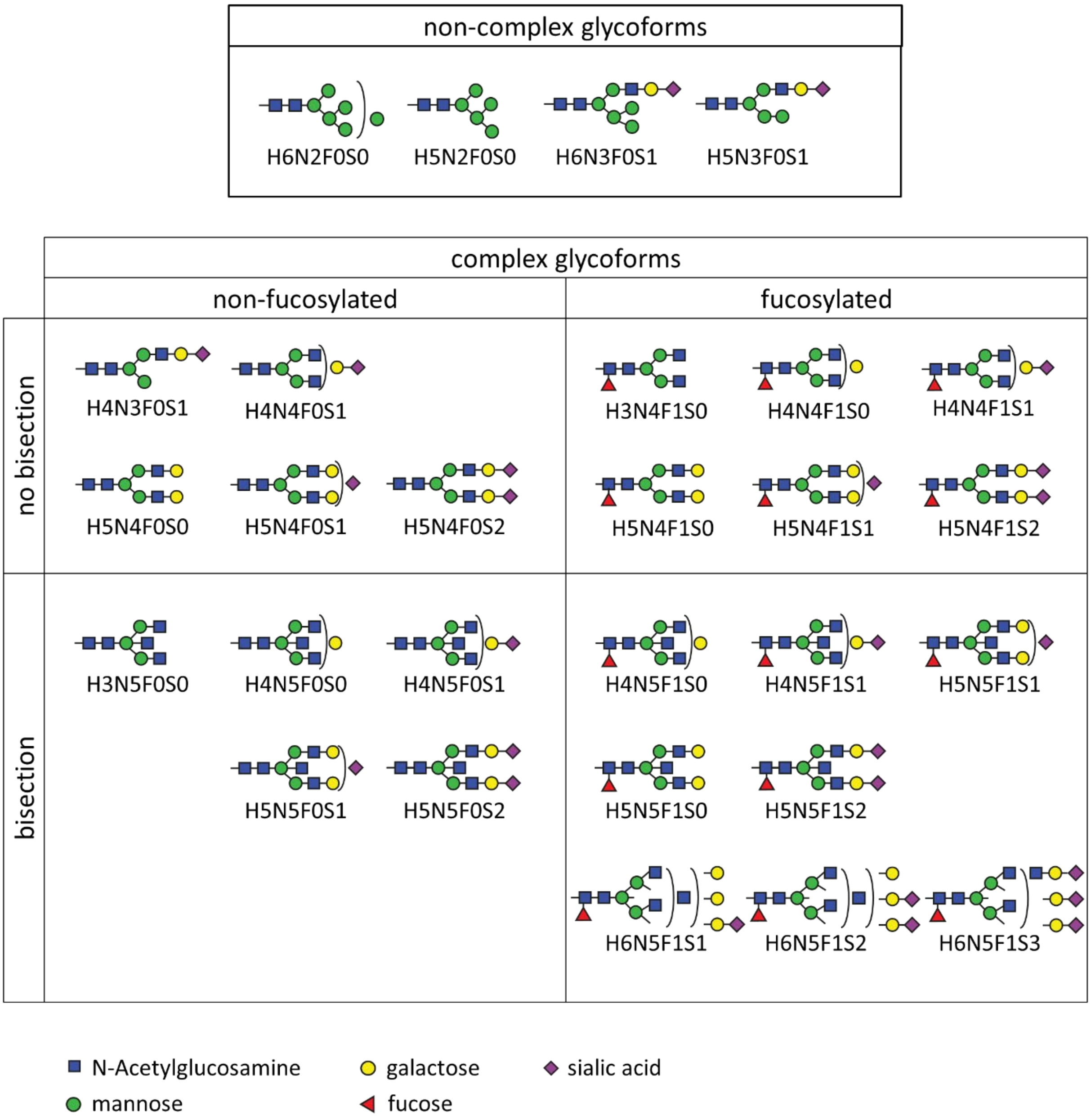


**Supplementary Fig. 2: Glycoform compositions found in IgA1, IgA2 and IgG.**

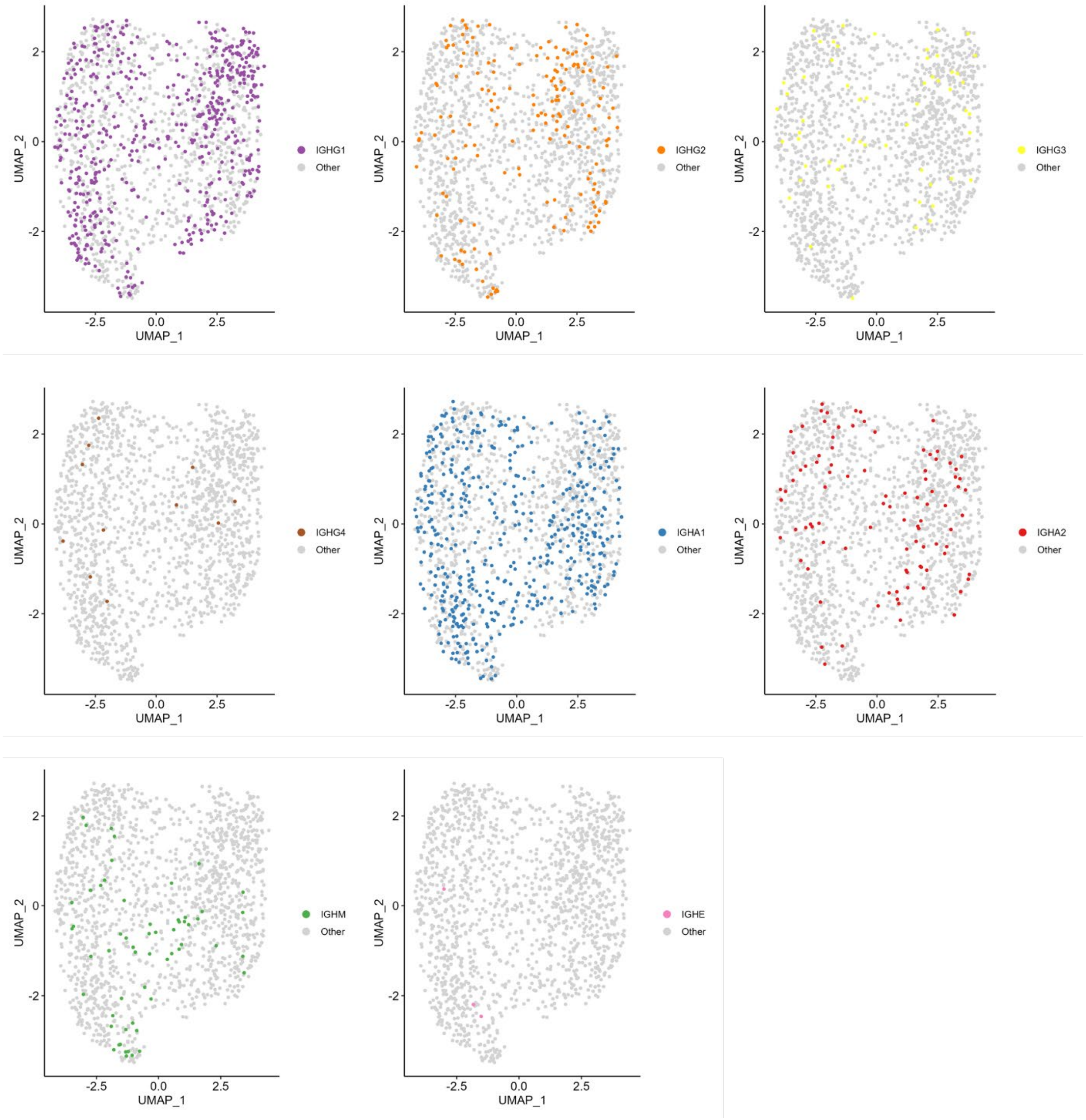


**Supplementary Fig. 3: Single cell RNA sequencing cell clustering of plasma cells sorted from bone marrow of healthy donors.** Shown are combined data from 3 donors visualized in UMAP colored by antibody subclass.

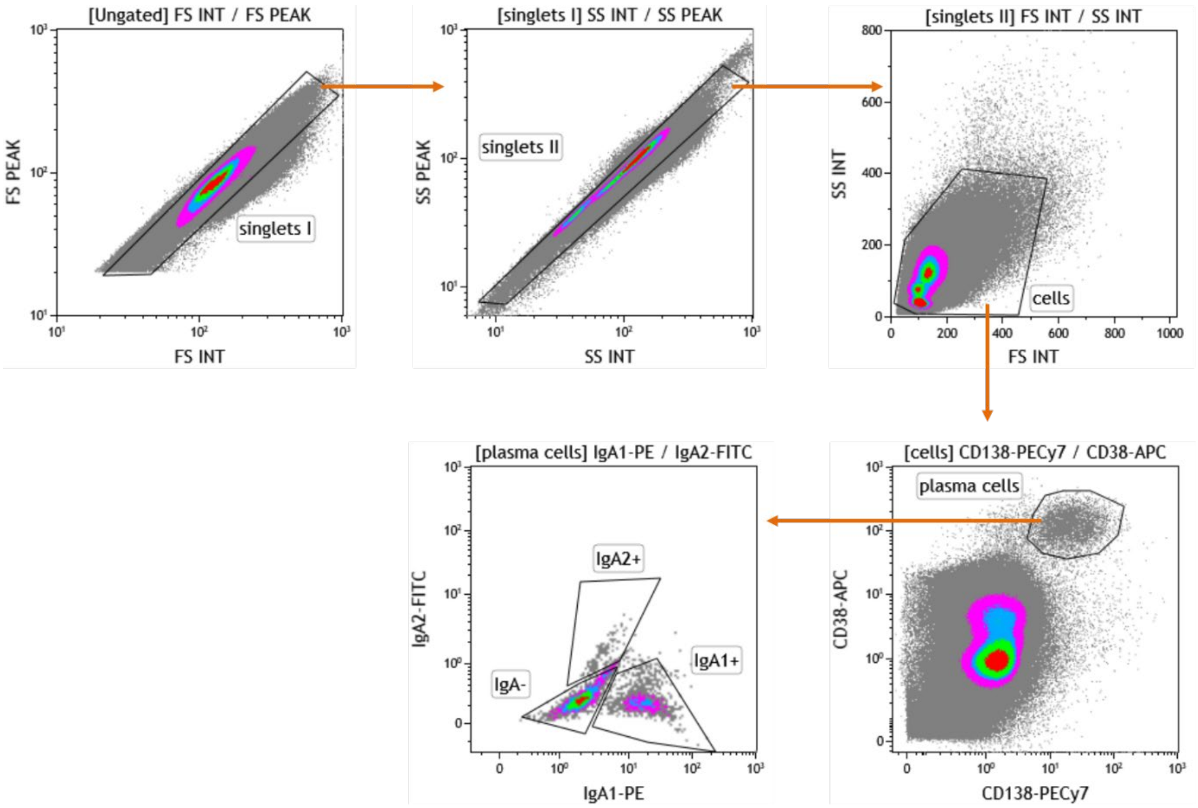


**Supplementary Fig. 4: Gating strategy for plasma cell evaluation.** First doublets were included by comparing forward scatter (FS) and sideward scatter (SS) peak against integral (INT). The remaining cells were analyzed for CD38 and CD138 expression and plasma cells gated as CD38$^{++}$CD138$^{++}$. Plasma cells were then analyzed for IgA1 and IgA2 positivity.

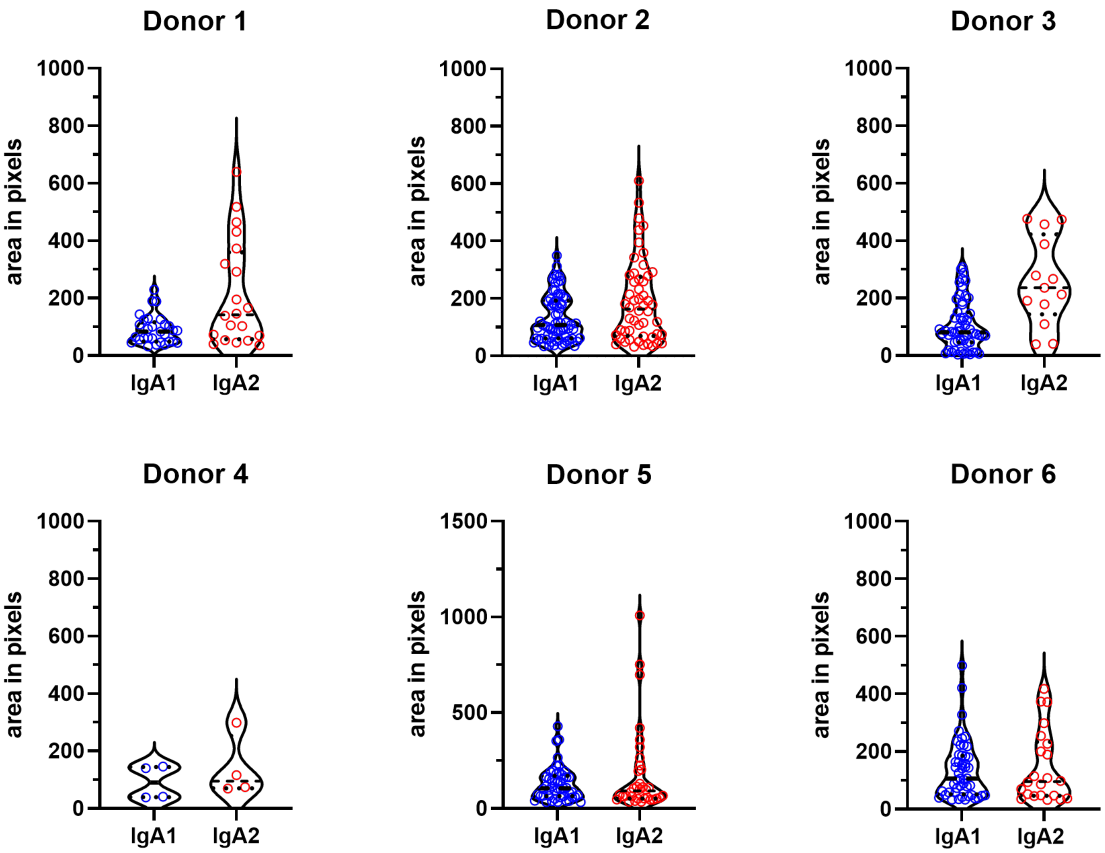


**Supplementary Fig. 5: Single spot sizes of the ELISpot assays.** ELISpots of human bone marrow cells using detection antibodies against IgA1 and IgA2. Shown are the violin blots for each individual donor.
